# Transforming chemigenetic bimolecular fluorescence complementation systems into chemical dimerizers using chemistry

**DOI:** 10.1101/2023.12.30.573644

**Authors:** Pratik Kumar, Alina Gutu, Amelia Waring, Timothy A. Brown, Luke D. Lavis, Alison G. Tebo

## Abstract

Chemigenetic tags are versatile labels for fluorescence microscopy that combine some of the advantages of genetically encoded tags with small molecule fluorophores. The Fluorescence Activating and absorbance Shifting Tags (FASTs) bind a series of highly fluorogenic and cell-permeable chromophores. Furthermore, FASTs can be used in complementation-based systems for detecting or inducing protein-protein interactions, depending on the exact FAST protein variant chosen. In this study, we systematically explore substitution patterns on FAST fluorogens and generate a series of fluorogens that bind to FAST variants, thereby activating their fluorescence. This effort led to the discovery of a novel fluorogen with superior properties, as well as a fluorogen that transforms splitFAST systems into a fluorogenic dimerizer, eliminating the need for additional protein engineering.

## Introduction

The use of fluorescence microscopy in biology has expanded thanks to new developments in microscope technology and advances in fluorescent labels such as fluorescent proteins and fluorophores^1^. While fluorescent proteins still dominate as markers for live-cell fluorescence microscopy,^2^ the introduction of hybrid small-molecule:protein (“chemigenetic”) tags has expanded the choice of labels available to researchers^3^. Chemigenetic systems combine a genetically encoded protein tag with a synthetic, small-molecule fluorophore.

Fluorescence Activating and Absorption Shifting Tags (FASTs) are an example of such systems. The original FAST was developed via the directed evolution of photoactive yellow protein (PYP) from *H. Halophila* to bind push–pull chromophores based on a hydroxybenzylidene rhodanine scaffold.^4^ These compounds are highly fluorogenic, enabling robust no-wash protocols. Subsequent work on the FAST protein yielded a more promiscuous version of FAST, pFAST,^5^ which binds a broader panel of fluorogens. Leveraging the sequence diversity in the parent PYP family yielded the promising new variant, RspAFAST that shows superior performance with first-generation fluorogens.^6^ FAST has also been developed as a bimolecular fluorescence complementation (BiFC) reporter for use in detecting protein:protein interactions (PPIs) by splitting the protein between residues 114 and 115.^7^ This split site was generalizable^8,9^ and could be applied to the second-generation RspAFAST protein; this system, known as splitFAST2,^6^ shows a lower non-specific background and a higher turn-on for known PPIs making it an attractive system for BiFC. A unique feature of splitFAST is its reversibility, allowing monitoring of protein association and dissociation. Applying the same split site to pFAST leveraged the increased inherent affinity of the two protein halves, transforming the split version of pFAST into a fluorogenic dimerizer system called CATCHFIRE that uses known ligands.^10^

In addition to optimizing chemigenetic systems using the power of protein engineering, these labeling systems can also be improved by employing organic chemistry on the small-molecule fluorescent ligand. FAST fluorogens consist of an electron-donating phenol moiety and an electron-withdrawing rhodanine headgroup; the structure–activity relationships of FAST fluorogens have been explored at both portions of the molecule.^5,11,12^ Adding electron-donating substituents to the phenol ring has proven to be a useful avenue for red-shifting the excitation and emission spectra of the molecules while retaining high fluorogenicity, similar to analogous substitutions made to fluorogens for RNA aptamers.^13^ Exploring different rhodanine headgroups and analogous structures has provided avenues for developing blue-shifted derivatives. Nevertheless, the promiscuity of the FAST system has made chemistry a secondary optimization tool that has mainly been employed to change spectral properties; a systematic exploration of structure–activity relationships has not been performed.

Here, we identify new FAST fluorogens with improved properties. We synthesized a comprehensive set of hydroxybenzylidene rhodanines with different methyl and methoxy substitution patterns on the phenolic moiety. We discovered that the mixed 3-methyl-5-methoxy derivative is a superior general-use fluorogen with pFAST and RspAFAST. We also discovered that the 2,5-dimethoxy derivative transforms a standard splitFAST system into a fluorogenic dimerizer without additional protein engineering. Together, these examples demonstrate that the modular FAST system can be further improved through the development of new ligands to monitor and elicit protein–protein interactions.

## Results and Discussion

The new fluorogens bridge the gap to explore both the nature and the position of the substituent on the phenol and how it affects the properties of the fluorogen. Substitutions at the previously identified third and fifth positions^11,12^ were varied with additional substitution at the second position. We also synthesized two unreported molecules, **3** and **5**. All the molecules **1**–**6** are capable of binding with the FASTs RspAFAST and pFAST (Table 1, S1). The affinities for these complexes range from sub-nanomolar to micromolar, depending on the protein variant and fluorogen. We further characterized these molecules for their spectral and photophysical properties. As with previously synthesized fluorogens, the novel molecules bind to FASTs with a characteristic red shift in absorption upon binding due to deprotonation of the fluorogen as well as an increase in quantum yield (Figure 1). Most fluorogens exhibit little variation in their spectra as a function of which FAST variant they are bound to. However, **6** exhibits a ∼25 nm red-shift in absorption when bound to RspAFAST versus pFAST (*λ*_max_ of 533 nm and 507 nm, respectively). This is likely due to specific hydrogen bonding interactions in the ligand binding pocket, as has been previously reported in other FAST variants.^8^

**Table 1.** Fluorogen properties with pFAST.

| Fluorogen | pK<sub>a</sub> | K<sub>D,app</sub><br>(nM) | τ<br>(ns) | λ<sub>max</sub><br>(nm) | λ<sub>em</sub><br>(nm) | φ<sup>a</sup> | ε (M<sup>-1</sup>cm<sup>-1</sup>) | Brightness |
| --- | --- | --- | --- | --- | --- | --- | --- | --- |
| **1** | 8.7 | 7.9<sup>b</sup> | 2 | 482 | 540 | 0.38 | 50500 | 19190 |
| **2** | 8.7 | 10<sup>b</sup> | 3.57 | 502 | 557 | 0.68 | 54600 | 37128 |
| **3** | 8.7 | 1 | 3.63 | 515 | 577 | 0.67 | 45000 | 30150 |
| **4** | 8.3 | 60<sup>b</sup> | 3.49 | 520 | 597 | 0.57 | 41800 | 23826 |
| **5** | 7.5 | 52 | 3.08 | 506 | 571 | 0.49 | 44500 | 21805 |
| **6** | 8 | 2 | 3.72 | 507 | 565 | 0.64 | 42900 | 27456 |

K<sub>D,app</sub>: Apparent binding affinity; τ: fluorescence lifetime; λ<sub>max</sub>: absorbance maximum;
λ<sub>em</sub>: fluorescence emission maximum; φ: fluorescence quantum yield
<sup>a</sup> absolute measurement <sup>b</sup> data taken from citation <sup>5</sup>

**Figure 1.**
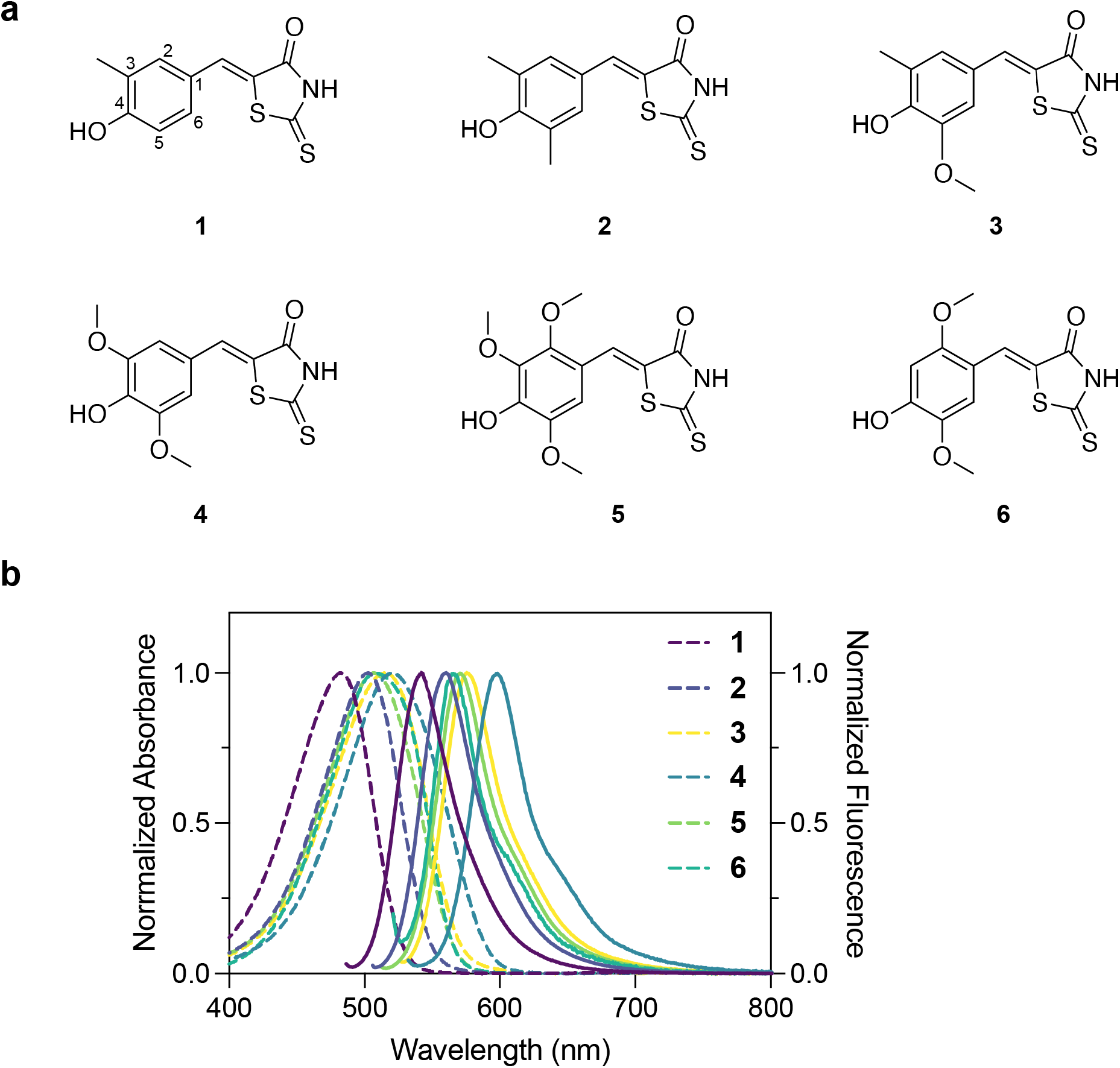
**a)** FAST fluorogen structures. **b)** Absorbance (dotted lines) and emission (solid line) spectra for pFAST:fluorogen complexes.

The overall brightest fluorogen is the previously reported **2**,^11,14^ however, the novel fluorogen **3** displays a similar fluorescence quantum yield and slightly lower molar extinction coefficient (45000 vs 54600 M^-1^cm^-1^). The fluorescence quantum yield of the fluorogens when bound to either RspAFAST or pFAST was measured and found to be equivalent between the two proteins. Similarly, the substitution position on the phenol ring did not appear to strongly influence the binding affinity of the fluorogens. For example, pFAST displays the tightest binding affinity for most of the previously published fluorogens, including the first-generation fluorogen **1** and the second-generation fluorogens, **2** and **4**. It also forms a tight complex with **3** and **6** with apparent binding affinities of 1 nM and 2 nM respectively. RspAFAST also binds the novel fluorogens well, with apparent binding affinities in the nanomolar range (Table S1). As might be expected due to increased steric bulk, the third methoxy substitution of **5** results in a ∼10-fold lower binding affinity.

To test the utility of this series of fluorogens, we evaluated their use in scanning confocal microscopy. We began by using a nuclear-localized construct, histone H2B-pFAST, and evaluated the relative brightness of each fluorogen in cells. The excitation power and wavelength were chosen to optimize excitation for each fluorogen while standardizing the illumination power. The overall brightest complex was H2B-pFAST:**2** while H2B-pFAST:**3** was the second brightest, consistent with its molar extinction coefficient and the tight affinity of pFAST for this molecule (Figure 2). We also tested the ability of pFAST to properly localize to the lumen of the mitochondria, the endoplasmic reticulum, the golgi body, and peroxisomes (Figure 2). We observed that pFAST was able to localize correctly to all subcellular locations, although for the peroxisome-targeted construct a proportion of cells showed diffuse cytosolic fluorescence. We quantified each localization with the series of fluorogens and found that **2** performed the best across the series of localizations, except for the mitochondria, where compound **3** was brighter (Figure S1). Nevertheless, the novel fluorogen **3** was useful as a general-purpose fluorogen, outperforming many of the first- and second-generation fluorogens.

**Figure 2.**
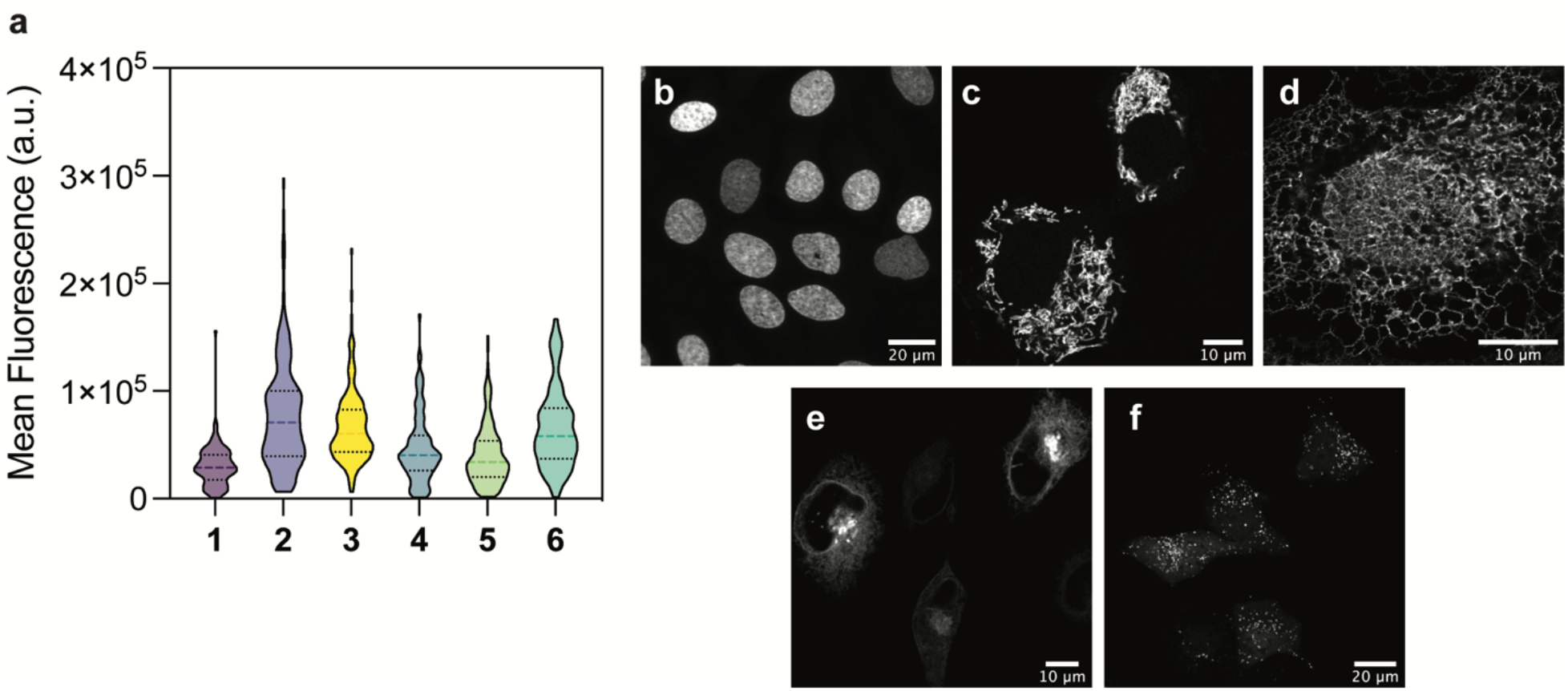
**a)** Live cell brightness of fluorogens when bound to H2B-pFAST. **b-f)** representative images of pFAST localized to the nucleus (**b**), mitochondrial lumen (**c**), endoplasmic reticulum lumen (**d**), Golgi body lumen (**e**), and interior of peroxisomes (**f**). All experiments were performed with 10 µM fluorogen.

As these new fluorogens show appropriate affinity and fluorogenic activity with full-length FASTs, we sought to evaluate their use as fluorogens with splitFAST. One of the key applications of FAST, where it excels beyond other available fluorescent proteins and tags, is its ability to be used as a reporter or inducer of molecular proximity as a so-called “split” system. To characterize the system, we fused each half of FAST with FRB or FKBP, which allows one to trigger molecular proximity by the addition of rapamycin, thus separating the effect of the fluorogen from the detection of molecular proximity. In the absence of rapamycin, the fluorescence from a given splitFAST:fluorogen combination arises due to spontaneous complementation driven by the affinity of the ternary complex comprised of the two fragments of splitFAST and the fluorogen.^6^ The addition of rapamycin then triggers the formation of a protein-protein interaction and forces the complementation of splitFAST, providing a read-out of maximal complementation.

We assessed the performance of four fluorogens, **2, 3, 5**, and **6**, with splitFAST2, a recently reported improved version of splitFAST.^6^ The N-terminal fragment (NFAST) was fused to FRB while the C-terminal fragment (CFAST) was fused to FKBP. Scanning line confocal microscopy was used to measure the appearance of splitFAST signal post-rapamycin addition with each fluorogen in a time course acquisition (Figure 3a-d). All tested fluorogens formed a fluorescent complex with splitFAST2 post-rapamycin addition. Interestingly, while fluorescence onset post-rapamycin addition was rapid with compounds **2, 3**, and **5**, cells treated with compound **6** were strongly fluorescent and largely unaffected by the addition of rapamycin, indicating fluorogen-dependent self-complementation (Figure 3d). Compounds **2** and **3** demonstrated a similar fluorescence-fold increase upon the addition of rapamycin, with a mean increase of ∼9-fold post-rapamycin. Compound **5** had a slightly elevated F/F_0_ (∼11-fold), however, exhibited lower overall brightness of the complex when mean fluorescence values are considered (Figures 3e and S2). The high self-complementation of **6** in the splitFAST association assay suggested that this molecule may behave as a fluorogenic dimerizer (Figure 3e). To examine this, we compared the interaction of **6** with splitFAST2 with the dimerizing system, CATCHFIRE, and its best-performing inducer, match_550_ (or, HBR-2,5DM) (Figure S3). The two compounds behaved similarly, with very little change in fluorescence before and after rapamycin addition. Compound **6** displayed a 1.3-fold increase in fluorescence, while CATCHFIRE displayed a 1.1-fold increase in fluorescence. Interestingly, both compound **6** and match_550_ display a 2,5-substitution pattern. Examination of a recent NMR structure^12^ and previous mutational analysis^14^ suggests that this substitution pattern may result in particularly high affinities due to hydrophobic interactions with the substituent at the 2 position, resulting in tight complex formation with the N-terminal fragment of FAST, which is key for the functioning of splitFAST^6^.

**Figure 3.**
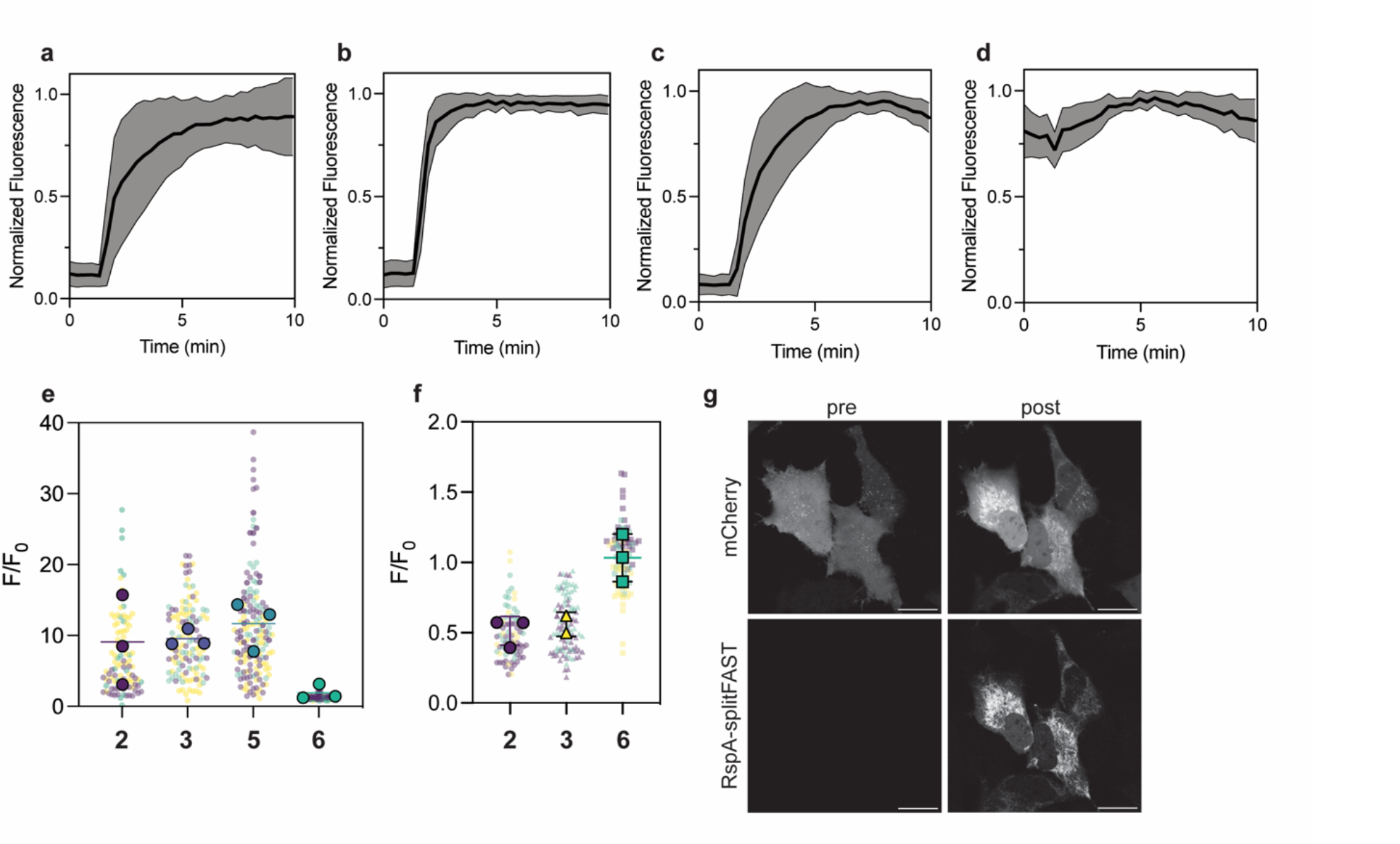
**(a-d)** Representative mean traces of splitFAST2 association with fluorogens **2 (a), 3 (b), 5 (c)**, and **6 (d)** labeled at 5 µM upon the addition of 500 nM rapamycin. **(e)** Summary data of fluorescence fold increase upon rapamycin addition. Small points are individual ROIs colored by experiment and large points are mean values across an experiment. **(f)** Summary of fluorescence fold decrease upon rapamycin addition for the dissociation of FKBP_F36M_ homodimer. **(g)** Representative images of recruitment of mCherry-CFAST to the mitochondrial membrane upon addition of 5 µM **6**.

To further evaluate the performance of **3** for use with splitFAST and of **6** as a dimerizer, we tested the ability of the splitFAST2:fluorogen complexes to dissociate (Figure 3f). To do this, we introduced the mutation F36M into FKBP, which results in the formation of a homodimer complex with micromolar affinity.^15^ In this case, the addition of rapamycin dissociates the complex. As in the association assay, **2** and **3** performed similarly, displaying a rapid decrease in fluorescence upon rapamycin addition (Figure S4). Conversely, **6** did not respond to the addition of rapamycin, which is consistent with it behaving as a fluorogenic dimerizer.

To further demonstrate the ability of **6** to function as a fluorogenic dimerizer, we used an assay that monitors the recruitment of fragments of splitFAST2 to the mitochondria. As in the previously reported CATCHFIRE system, the N-terminal fragment was targeted to the outer membrane of the mitochondria, while the C-terminal fragment was fused to cytosolic mCherry. The localization of mCherry-RspA-CFAST as well as the signal from splitFAST2 was monitored over time (Figure 3g). Analogously to previous reports with CATCHFIRE, the addition of **6** drove the recruitment of mCherry-RspA-CFAST to the mitochondria. This observation is consistent with the association and dissociation assays that **6** can act as a fluorogenic dimerizer.

## Conclusion

Here, we describe the development of new fluorogens for FAST tags that perform well in cellular assays. Compound **3**, identified through exploration of the substituents on the phenol ring of hydroxybenzylidene rhodanine derivatives forms a bright complex with various FAST variants. With the newly identified RspAFAST, **3** forms a complex with comparable brightness as the previously reported **2**. Compound **3** is also compatible with the recently engineered pFAST for labeling the lumen of various cellular organelles.

We have also described the use of **6** as a fluorogenic dimerizer. In contrast to previous reports,^10^ this molecule functions well without the need to change the protein sequences used. This additional experimental flexibility allows one to use the protein identified as the best-performing reporter for molecular proximity as a fluorogenic dimerizing system simply by changing the identity of the small molecule used. The resulting complex is bright and amenable to live imaging. Furthermore, one can likely exert additional control over the system by tuning the amount of **6** used. We expect this additional experimental versatility to enable both the monitoring and the perturbation of molecular proximity and protein-protein interactions.

## Acknowledgements

The authors would like to thank Dr. Chris Obara (Janelia Research Campus) for this gift of plasmids encoding the sequences for ER and Golgi body localization and the Tool Translation Team (Janelia Research Campus) for research support and feedback.

