## Supplementary Information for "Transforming chemigenetic bimolecular fluorescence complementation systems into chemical dimerizers using chemistry"

#### Supplemental Data

Table S1. Fluorogen properties with RspAFAST

| Fluorogen | $K_D$ (nM) | $\lambda_{max}$ (nm) | $\lambda_{em}$ (nm) | $\varphi^1$ |
|---|---|---|---|---|
| **1** | $31^2$ | 482 | 544 | 0.34 |
| **2** | $11^2$ | 503 | 559 | 0.68 |
| **3** | 20 | 521 | 574 | 0.65 |
| **4** | $74^2$ | 536 | 598 | 0.5 |
| **5** | 240 | 515 | 571 | 0.5 |
| **6** | 8 | 533 | 568 | 0.65 |

$^1$abolute measurement $^2$ from ref [1]

Table S2. Fluorogen properties in solution

| | 0.1N HCl | | PBS | | 0.1N NaOH | |
|---|---|---|---|---|---|---|
| Fluorogen | $\lambda_{max}$ (nm) | ε ($M^{-1}cm^{-1}$) | $\lambda_{max}$ (nm) | ε ($M^{-1}cm^{-1}$) | $\lambda_{max}$ (nm) | ε ($M^{-1}cm^{-1}$) |
| **1** | 403 | 32900 | 401 | 31400 | 461 | 38700 |
| **2** | 394 | 9500 | 401 | 31700 | 474 | 40700 |
| **3** | 400 | 19000 | 407 | 30700 | 482 | 38700 |
| **4** | 411 | 29700 | 408 | 26100 | 487 | 34600 |
| **5** | 412 | 26600 | 418 | 17400 | 480 | 33500 |
| **6** | 422 | 11700 | 442 | 19000 | 492 | 35900 |

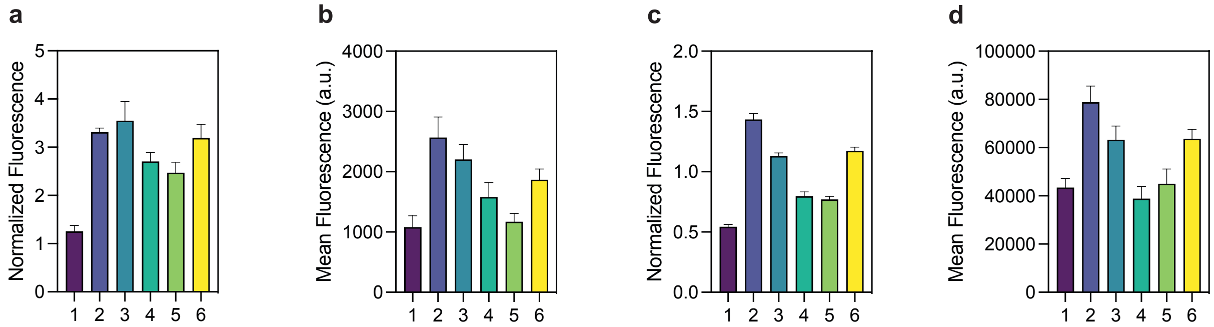


**Figure S1.** Brightness comparison for all fluorogens in the lumen of the (**a**) mitochondria, normalized to iRFP signal, (**b**) golgi body, (**c**) endoplasmic reticulum, normalized to iRFP signal, and (**d**) peroxisomes. All fluorogens were used at 10 μM concentration.

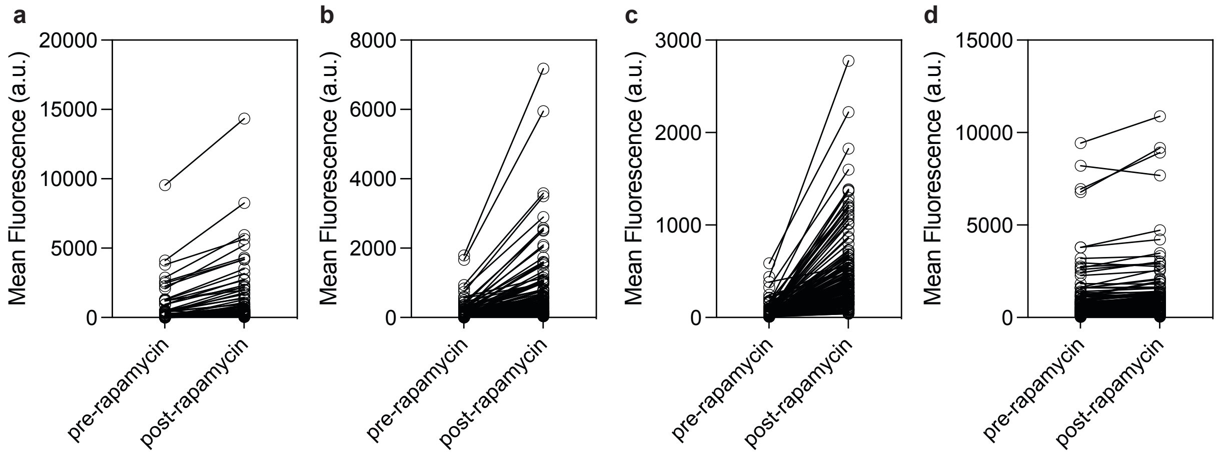


**Figure S2.** Mean fluorescence values pre- and post-rapamycin addition for **2** (**a**), **3** (**b**), **5** (**c**), and **6** (**d**). All data from all experiments are plotted.

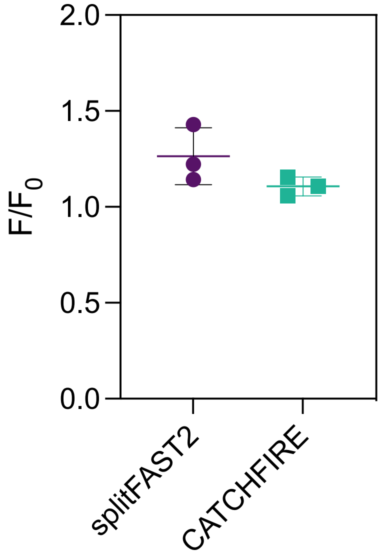


**Figure S3.** Comparison of the association of splitFAST2:**6** with CATCHFIRE in the presence of 5 μM fluorogen upon addition of 500 nM rapamycin. The mean $F/F_0$ from each of the three experiments is plotted (n = 82 and 61 cells).

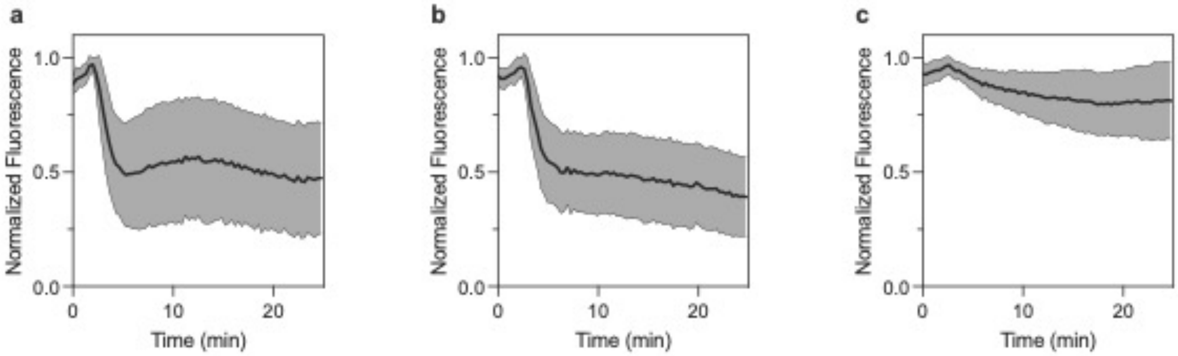


**Figure S4.** Representative time course data of splitFAST2 dissociation labeled with 5 μM fluorogen upon the addition of 2.5 μM rapamycin. **(a)** fluorogen **2**, **(b)** fluorogen **3**, **(c)** fluorogen **6.**

#### Material and methods

##### Organic Synthesis

**General.** Commercial reagents were the highest quality available and used as received. Solvents for reactions were of anhydrous grade, purchased in septum-sealed bottles, and stored under an inert atmosphere. Reactions were conducted in round-bottomed flasks or septum-sealed crimp-top microwave reaction vials (Biotage) containing Teflon-coated magnetic stir bars. All reactions were conducted under an inert atmosphere of Ar(g) and protected from light using Al foil unless otherwise noted. Heating of reaction mixtures was achieved through aluminum blocks on top of a stirring hotplate equipped with an electronic contact thermometer. Microwave reactions were conducted using a commercial microwave reactor (Biotage initiatior+). Reactions were monitored either by thin layer chromatography (TLC) on precoated TLC glass plates (silica gel 60 F254, 250 μm thickness) or by tandem liquid chromatography–mass spectrometry (LC–MS; Shimadzu LCMS 2020, Phenomenex Kinetex 30 × 2.1 mm 2.6 μm C18 column, 1–10 μL injection, 5–98% $CH_3CN/H_2O$ linear gradient with constant 0.1% v/v $HCO_2H$, 6 min run, 1 mL/min flowrate, ESI, positive ion mode). TLC plates were visualized either by UV illumination or by developing the TLC with ceric ammonium molybdate or $KMnO_4$.

Reaction products were purified either by flash chromatography on Biotage Isolera automated purification system using prepacked silica gel columns and/or by preparative high-pressure liquid chromatography (HPLC; Agilent 1200, Phenomenex Gemini–NX 150 × 30 mm 10 μm C18 110 Å column, 42 mL/min flowrate) under the indicated solvent gradient conditions. Analytical HPLC analyses were performed on a LC-MS system (Agilent 1200, Phenomenex Gemini–NX 150 × 4.6 mm 5 μm C18 110 Å column, 1 mL/min flowrate) or an analytical HPLC (Shimadzu UFLC, Phenomenex Gemini–NX 150 × 4.6 mm 5 μm C18 110 Å column, 1 mL/min flowrate) under the indicated conditions. All the HPLC systems are fitted with a diode array detector. High-resolution mass spectrometry was obtained from the High Resolution Mass Spectrometry Facility at the University of Iowa. NMR spectra were recorded on Bruker Avance 400 MHz spectrometer and processed through MestReNova. Deuterated solvents were used as purchased. $^1H$ and $^{13}C$ chemical shifts (δ) were referenced to TMS or residual solvent peaks. $^{19}F$ chemical shifts were referenced to $CFCl_3$. Data for $^1H$ NMR spectra (Appendix I) are reported as follows: chemical shift (δ ppm), multiplicity (s = singlet, d = doublet, t = triplet, q = quartet, p = pentet (quintet), dd = doublet of doublets, dt = doublet of triplets, m = multiplet, br = broad signal), coupling constant (Hz), and integration. Data for $^{13}C$ NMR spectra (Appendix I) are reported by chemical shift (δ ppm) with hydrogen multiplicity (C, CH, $CH_2$, $CH_3$) information obtained from DEPT spectra.

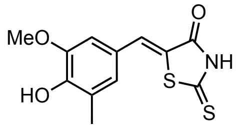


**5-(4-hydroxy-3-methoxy-5-methylbenzylidene)-2-thioxothiazolidin-4-one (3):** A suspension of 3-methyl-4-hydroxy-5-methoxybenzaldehyde (249.5 mg, 1.5 mmol, 1 equiv) and rhodanine (200 mg, 1.5 mmol, 1 equiv) in water (10 mL) was placed in a sealed vial and heated using a microwave system for 24 h at 90 ºC. The reaction was cooled, and the resulting yellow precipitate was collected by filtration, washed with water (~50 mL), and dried under high vacuum to obtain **3** as a yellow solid (305 mg, 72.2%). $^{1}$H NMR (400 MHz, DMSO-d$_6$) δ 13.69 (s, 1H), 9.59 (d, *J* = 2.2 Hz, 1H), 7.52 (t, *J* = 2.6 Hz, 1H), 7.03 (t, *J* = 2.7 Hz, 1H), 6.98 (t, *J* = 2.5 Hz, 1H), 3.86 (s, 3H), 2.17 (s, 3H). $^{13}$C NMR (101 MHz, DMSO-d$_6$) δ 195.50 (C), 169.43 (C), 147.83 (C), 147.56 (C), 132.96 (CH), 126.47 (CH), 125.42 (C), 123.59 (C), 120.94 (C), 111.90 (CH), 55.92 (CH$_3$), 15.85 (CH$_3$). $^{19}$F NMR (376 MHz, DMSO-d$_6$) δ -61.58. HRMS (ESI) calculated for $C_{12}H_{10}NO_3S_2$ [M-H]$^-$ = 280.0108, found 280.0107.

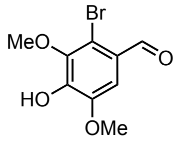


**2-bromo-4-hydroxy-3,5-dimethoxybenzaldehyde (S1)**[1]**:** Syringaldehyde (5 g, 27.45 mmol, 1 equiv) was dissolved in $CH_2Cl_2$ (475 mL). $Br_2$ (2.1 mL, 6.58 g, 41.17 mmol, 1.5 equiv) was added dropwise. This red solution was stirred for 16 h at ambient temperature. The reaction mixture was concentrated under reduced pressure and the title compound was purified by crystallization from 1:1 v/v cyclohexane/EtOAc (300 mL) as off-white crystals (5.55 g, 77.5%).

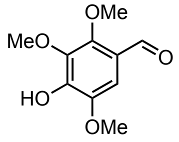


**4-hydroxy-2,3,5-trimethoxybenzaldehyde (S2)**[2]**: S1** (50 mg, 191 µmol, 1 equiv), $K_2CO_3$ (79.4 mg, 574 µmol, 3 equiv), and $CuCl_2$ (5 mg) was suspended in methanol (10 mL). The reaction vessel was sealed and the suspension was stirred for 24 h at 130 ºC. The reaction mixture was diluted with 6 N HCl and extracted with $CH_2Cl_2$. The combined organics were dried over $Na_2SO_4$ and concentrated under reduced pressure. Purification by $SiO_2$ gel chromatography (25 g $SiO_2$ column, 0–50% v/v EtOAc/hexanes, linear gradient) provided **S2** (25 mg, 61.6%).

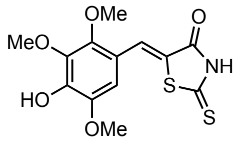

**5-(4-hydroxy-2,3,5-trimethoxybenzylidene)-2-thioxothiazolidin-4-one (5):** Compound **S2** (140 mg, 660 µmoles, 1.0 equiv) and rhodanine (88 mg, 660 µmoles, 1 equiv) was suspended in ethanol (10 mL). Piperidine (65 µL, 660 µmoles, 1 equiv) was added and the reaction mixture was stirred for 40 h at ambient temperature. The reaction mixture was acidified to pH ≈ 1 using 1 N HCl (25 mL). The resulting precipitate was collected by filtration, washed with water, and dried under high vacuum to obtain **5** as a yellow-orange solid (146.3 mg, 66.5%). $^1$H NMR (400 MHz, DMSO-$d_6$) δ 13.72 (s, 1H), 9.94 (s, 1H), 7.69 (s, 1H), 6.66 (s, 1H), 3.83 (s, 3H), 3.80 (s, 3H), 3.78 (s, 3H). $^{13}$C NMR (101 MHz, DMSO-$d_6$) δ 195.71 (C), 169.50 (C), 148.65 (C), 145.38 (C), 144.70 (C), 141.44 (C), 127.21 (CH), 122.24 (C), 115.75 (C), 106.27 (CH), 61.96 ($CH_3$), 60.43 ($CH_3$), 56.10 ($CH_3$). HRMS (ESI) calculated for $C_{13}H_{12}NO_5S_2$ $[M-H]^-$ = 326.0162, found 326.0162.

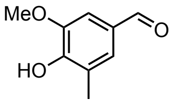


**4-hydroxy-3-methoxy-5-methylbenzaldehyde (S3)**[3]**:** 3-((dimethylamino)methyl)-4-hydroxy-5-methoxybenzaldehyde (4 g, 19.1 mmol, 1 equiv) was dissolved in acetic anhydride (20 mL). The resulting solution was heated to reflux for 18 h. The reaction mixture was concentrated under reduced pressure and the crude material was stirred in concentrated HCl (20 mL) for 2.5 h at ambient temperature. The resulting suspension was diluted with dioxane (30 mL). $SnCl_2$ (10.9 g, 57.4 mmol, 3 equiv) was added and the resulting black suspension was refluxed for 30 mins. The reaction mixture was cooled, diluted with water (50 mL), acidified with concentrated HCl (8 mL), and extracted with $CH_2Cl_2$ (25 × 4 mL). The combined organic layers were dried over $MgSO_4$ and concentrated under reduced pressure. Purification by $SiO_2$ gel chromatography (100 g $SiO_2$ column, 0–40% v/v EtOAc/hexanes, linear gradient) provided **S3** as a white solid (2.5 g, 65.3% over three steps).

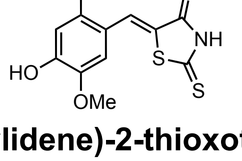


**5-(4-hydroxy-2,5-dimethoxybenzylidene)-2-thioxothiazolidin-4-one (6):** A suspension of 2,5-dimethoxy-4-hydroxy-benzaldehyde (1 g, 5.5 mmol, 1 equiv) and rhodanine (877 mg, 6.6 mmol, 1.2 equiv) in water (10 mL) was placed in a sealed vial and heated using a microwave system for 24 h at 90 ºC. The yellow precipitate was collected by filtration, suspended in ethanol (60 mL), and stirred for 16 h at ambient temperature. The resulting solid was collected by filtration and dried under high vacuum to obtain **6** as an orange solid (1.23 g, 75.4%). $^1$H NMR (400 MHz, DMSO-$d_6$) δ 13.61 (s, 1H), 10.32 (s, 1H), 7.75 (s, 1H), 6.84 (s, 1H), 6.60 (s, 1H), 3.81 (s, 3H), 3.79 (s, 3H). $^{13}$C NMR (101 MHz, DMSO-$d_6$) δ 195.55 (C), 169.55 (C), 154.87 (C), 152.30 (C), 142.33 (C), 127.68

(CH), 120.11 (C), 112.83 (CH), 111.59 (C), 100.46 (CH), 56.16 ($CH_3$), 55.98 ($CH_3$). HRMS (ESI) calculated for $C_{12}H_{12}NO_4S_2$ $[M-H]^-$ = 296.0057**,** found 296.0056.

#### Molecular and Cell Biology

##### Expression vectors

The construct for targeting pFAST to the nucleus was ordered from Genscript for cloning into their pcDNA3.1+(Zeo). Constructs for targeting pFAST to organelles were designed and ordered as gBlocks from IDT and cloned downstream of the CMV promoter in plasmid Addgene #70113 via NEBuilder HiFi DNA Assembly kit (Biolabs). pFAST ORF (Addgene #172862) was fused to specific targeting sequences for luminal expression in various organelles. The Cox8 sequence was used for mitochondrial targeting (Addgene #55102); the tandem N- terminus Prolactin signal sequence/C- terminus KDEL for ER targeting (*gift from Dr. Chris Obara*); the ST6GAL1 signal sequence (*gift from Dr. Chris Obara*) for Golgi localization; and the C-terminus SKL (Addgene #54520) for peroxisome localization. For co-expression of both FAST and iRFP670 from the same promoter, IRES sequence was inserted between the two ORFs. iRF670 was fused to various sequences for extraluminal expression in organelles, such as Tomm 20 (Addgene #55146) for mitochondrial localization; Calnexin (Addgene #55891) for ER localization; Sys1 (Addgene #175092) for Golgi, and PXMP2 for peroxisomes (Addgene #55910). For nuclear pFAST localization, a stable U2OS cell line created in-house was used (*see below*). Sequences corresponding to mCherry-RspA-CFAST and tom20-CFP-RspA-NFAST were ordered from Twist cloned into the pTwist-CMV vector.

All mammalian expression plasmids generated in this study were deposited to Addgene (in progress).

##### Mammalian cell culture

U2OS cells (ATCC), HEK 293 (ATCC), and HeLA (ATCC) were cultured in Dulbecco's modified Eagle medium (DMEM, phenol red-free; # 17205CV Corning) supplemented with 10% (v/v) fetal bovine serum (FBS, #26140095 Gibco), 2 mM L-Glutamine (# 35050061 Gibco) and maintained at 37 °C in a humidified 5% (v/v) $CO_2$ setting. For routine maintenance, cells were passaged at 70-80% confluency by trypsinization. To generate a stable U2OS cell line, 750 ng of pcDNA3.1-H2B-pFAST was electroporated into 0.5e6 cells. After two days, 500 µg/mL zeocin was added. Cells were passaged three times and then sorted for medium levels of positive FAST expression. For other subcellular targets, U2OS or HeLA cells were transiently transfected using nucleofection (Lonza) with 1µg of each expression vector. For splitFAST imaging experiments HEK 293 or HeLa cells were transiently transfected using FuGENE (Promega) with 1 µg total DNA per µdish.

**Cell imaging and analysis**

Transiently transfected cells were imaged 20-48 hours post-transfection depending on the construct. Prior imaging, the transfected cells or stable H2B-FAST U2OS cells were seeded in either Nunc Lab-Tek (#155409) or Ibidi (#80826) 4-or 8-well slide chambers at a density that allows good coverage for imaging. When dual expression constructs were used, the media was supplemented with 25 µM biliverdin 4-8 hours after seeding. Before imaging fresh FBS-free media was mixed with 10 µM FAST fluorogens and added to cells. Imaging was done on a Leica SP8 Falcon or Leica Stellaris 8 equipped with an environmental chamber at 37 °C, 5% $CO_2$ air. Images were acquired as Z-stacks at optimal settings (avoidance of detector saturation, minimizing photobleaching, and minimize cross-excitation or emission bleed-through). To account for different excitation and emission maxima of each compound, we adjusted accordingly the excitation line (Table 1) and detection window (+/-20 nm) while maintaining the laser power constant. For analysis of nuclei in stable H2B-FAST U2OS, prior to FAST fluorogens addition, 6 µg/ml Hoechst dye was used (#3342 ThermoFisher) by staining for 15 min at 37 °C. All images were first background subtracted in Image J. The Hoechst channel (405 nm excitation, 415-500 nm emission) was used to create binary images by: making a sum projection, gaussian blurring, Huang automatic thresholding in Image J. The nuclei objects generated with the Hoechst channel were used to extract normalized integrated density of grayscale values in the FAST channel. For quantification of FAST signal of mitochondria, ER, or Golgi, the iRFP670 channel (ex. 633 nm, emission 650 nm – 780 nm) was first used to create the binary images as was done with Hoechst channel for nuclei. To account for variation in expression relative to iRFP signal, the FAST signal for mitochondria and ER was divided to the iRFP signal of each object. For quantification of peroxisome signal, Imaris 9.7 (Bitplane) was used to generate surface models (with background subtraction, seed size 0.5 µm) in the iRFP670 channel and group peroxisomes on a per cell basis. Sum of FAST channel intensities in each object was extracted for quantification. To confirm localization of FAST in mitochondria, ER, and Golgi we also used commercial trackers for each organelle and labeled according to recommended instructions of each manufacturer. For mitochondria, we used 50 nM MitoTracker Deep Red FM (#M22426 ThermoFisher) for 30 min at 37 °C; washing then imaging (ex 633, emission 650-750 nm); for ER, we used 1 µM ER-Tracker Red (#E34250 ThermoFisher) for 30 min at 37 °C (ex 587, emission 625-660 nm); and for Golgi, 1x Golgi-ID Green detection reagent and Hoechst 33342 combo (#EnZ-51028 Enzo) for 30 min at 4 °C and 30 min at 37 °C.

For splitFAST imaging, HEK cells were seeded into ibidi µdishes coated with poly-D-lysine and reverse transfected with 1 µg DNA using FuGENE (Promega). After 48 hours, cells were imaged on a Leica Stellaris 8 equipped with a white light laser (440 nm – 750 nm). For imaging cells were exchanged into DMEM with HEPES, supplemented with 5 µM of the appropriate fluorogen, except for mitochondrial recruitment studies where DMEM only was used. For association assays, images were acquired every 20 seconds for ten minutes; rapamycin was added after the fifth image to a final concentration of 500 nM. For dissociation assays, images were acquired every 20 seconds for 15 minutes; rapamycin was added after the fifth image to a final concentration of 2.5 µM. For mitochondrial recruitment assays, the acquisition was begun and 10 µM fluorogen was added to give a final concentration of 5 µM. Cells from either the last (association) or first (dissociation) were segmented using cellpose 2.0[4]. A custom macro was written to use the cellpose segmentations in Fiji to produce masks and ROIs that were then used to automatically extract the mean fluorescence from tiff stacks. The resulting fluorescence data was cleaned and processed using a python script and then plotted in Prism 10 (GraphPad).

**Fluorescence lifetime imaging**

Fluorescence Lifetime Imaging was performed using Leica SP8 Falcon system and the LaSX 3.5.6 software for image acquisition and analysis. Stable H2B-FAST U2OS cells grow into 24-well glass bottom plate (# P24.1.5 H-N Cellvis) were incubated with 10 µM of FAST fluorogens in FBS-free media. Images were acquired with the 40X oil immersion 1.3 NA objective. Excitation (tuned at 20 MHz) and emission was achieved as described above. The acquired FLIM images were fitted with two component exponential function and the average mean $\tau$ of the amplitude-weighted lifetime was calculated (n=4).

**pFAST and RspAFAST protein purification**

Bacterial pET28a expression vectors expressing N-(6xHis)-pFAST and RspA protein were transfected into T7-Expresss competent cells (NEB), grown in Terrific Broth (#22-7711-022 Invitrogen) and induced for 4 h with 0.1 - 0.2 mM IPTG (# 15529-019 Invitrogen) at 37 °C with 220 RPM shaking. B-Per bacterial protein extraction reagent (#78248 ThermoFisher) supplemented with 1 mg/ml lysozyme (#12650-88-3 Sigma) and Pierce Universal Nuclease (#88702) was used to lyse the cells. The soluble 6×His- pFAST and 6×His-RspA proteins was affinity purified using Profinity IMAC Resin (#156-0137 Biorad) with a 0–200 mM imidazole elution gradient in 50 mM sodium phosphate buffer, 0.3M NaCl pH 8. The fractions containing the protein was pooled, concentrated by a Amicon Ultra-15 3 KDA cutoff spin concentrator (#900396, Millipore) with buffer exchange into 50 mM sodium phosphate, 0.15 M NaCl pH 7.4 (storage buffer) to remove the imidazole. Protein purity was assessed by analysis of Coomassie-stained PAGE-SDS gels (#NP0322 and # NP0002 ThermoFisher).

**Apparent binding affinities**

Apparent dissociation constants for pFAST and RspAFAST were determined as previously reported.[5,6] Titrations were carried out in technical triplicate with 20 or 50 nM protein and measured on a Biotek Cytation 5. The resulting curves were fit directly in Prism 10 using a one-site specific binding model.

**1-Photon spectroscopy of fluorogens at different pH**

Compounds **1-6** were prepared as stock solutions in DMSO and diluted such that the final DMSO concentration did not exceed 1% v/v. All measurements were taken at ambient temperature (22 ± 2 °C). Absorption spectra were recorded on a Cary Model 100 spectrometer (Agilent) using 1-cm path length 1.0-mL quartz microcuvettes from Starna Cells. Fluorogens **1-6** (10 μM) were dissolved in 0.1 N HCl (pH 1.1) or 1× PBS (pH 7.4) or 0.1 N NaOH (pH 12.8), and the resulting mixture was incubated for 1 h before recording their absorbance. Reported values for extinction coefficient ($\varepsilon$) are averages of at least three measurements.

**1-Photon spectroscopy of fluorogens with pFAST protein**

Compounds **1-6** were prepared as stock solutions in DMSO and diluted such that the final DMSO concentration did not exceed 1% v/v. All measurements were taken at ambient temperature (22 ± 2 °C). Absorption spectra were recorded on a Cary Model 100 spectrometer (Agilent) using 1-cm path length 1.0-mL quartz microcuvettes from Starna Cells. Fluorescence spectra were recorded on a Cary Eclipse fluorometer (Varian) using 1-cm path length 3.5-mL quartz cuvettes (Starna Cells). Fluorogens **1-6** (5 μM) and pFAST (10 μM) were dissolved in 1× PBS, pH 7.4, and the resulting mixture was incubated for 1 h before recording their absorbance or fluorescence. To measure the fold-increase of absorbance or fluorescence upon pFAST binding, a "no pFAST" control experiment was performed where an equivalent volume of PBS blank was added in place of the protein. Reported values for extinction coefficient ($\varepsilon$) are averages of at least two measurements.

**1-Photon spectroscopy of fluorogens with RspAFAST protein**

Compounds **1-6** were prepared as stock solutions in DMSO and diluted such that the final DMSO concentration did not exceed 1% v/v. All measurements were taken at ambient temperature (22 ± 2 °C). Absorption and fluorescence spectra were recorded on Cytation5 plate reader (Biotek) in a 96-well plate. Fluorogens **1-6** (1 μM) and RspAFAST (15 μM) were dissolved in 1× PBS, pH 7.4, and the resulting mixture was incubated for 1 h before recording their absorbance or fluorescence. To measure the fold-increase of absorbance or fluorescence upon pFAST binding, a "no pFAST" control experiment was performed where an equivalent volume of PBS blank was added in place of the protein. Reported values for extinction coefficient ($\varepsilon$) are averages of at least two measurements.

**Quantum yield determination**

All reported absolute fluorescence quantum yield values ($\Phi_f$) were measured under identical conditions using a Quantaurus-QY spectrometer (model C11374, Hamamatsu). This instrument uses an integrating sphere to determine photons absorbed and emitted by a sample. Measurements were performed using dilute samples (absorbance < 0.1), and self-absorption corrections were performed using the instrument software. Reported values are averages of at least two measurements.

### Appendix I

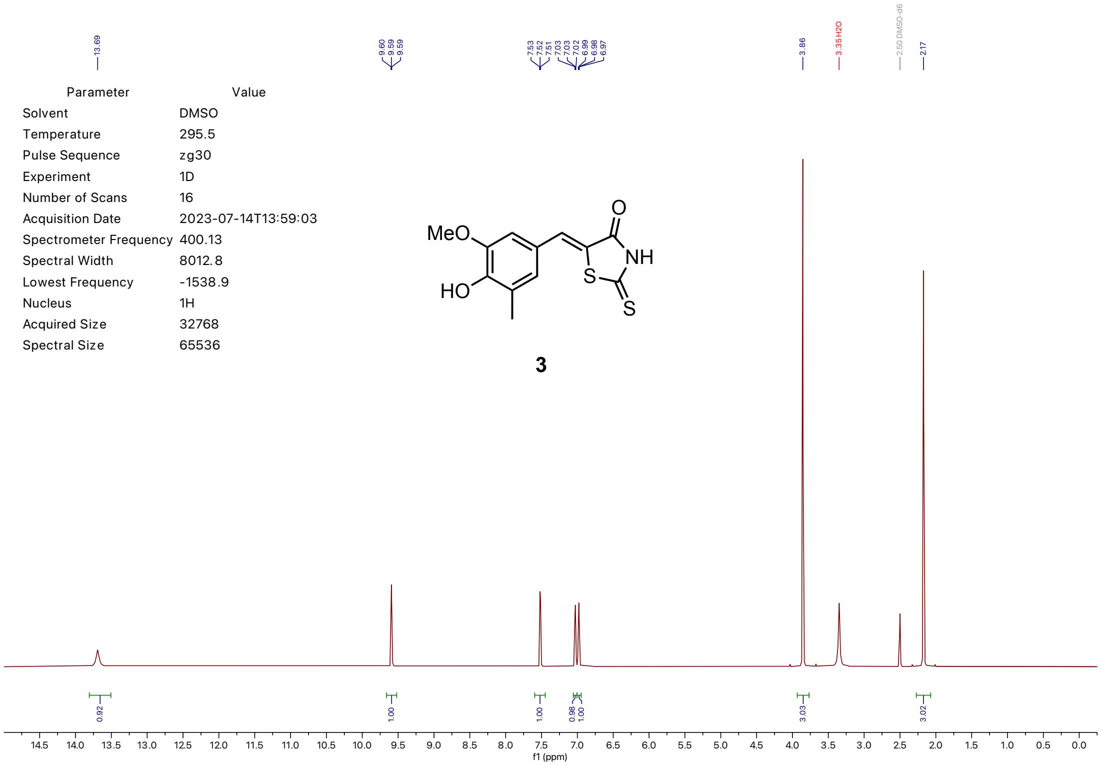


| Parameter | Value |
|---|---|
| Solvent | DMSO |
| Temperature | 295.5 |
| Pulse Sequence | zg30 |
| Experiment | 1D |
| Number of Scans | 16 |
| Acquisition Date | 2023-07-14T13:59:03 |
| Spectrometer Frequency | 400.13 |
| Spectral Width | 8012.8 |
| Lowest Frequency | -1538.9 |
| Nucleus | 1H |
| Acquired Size | 32768 |
| Spectral Size | 65536 |

**3**

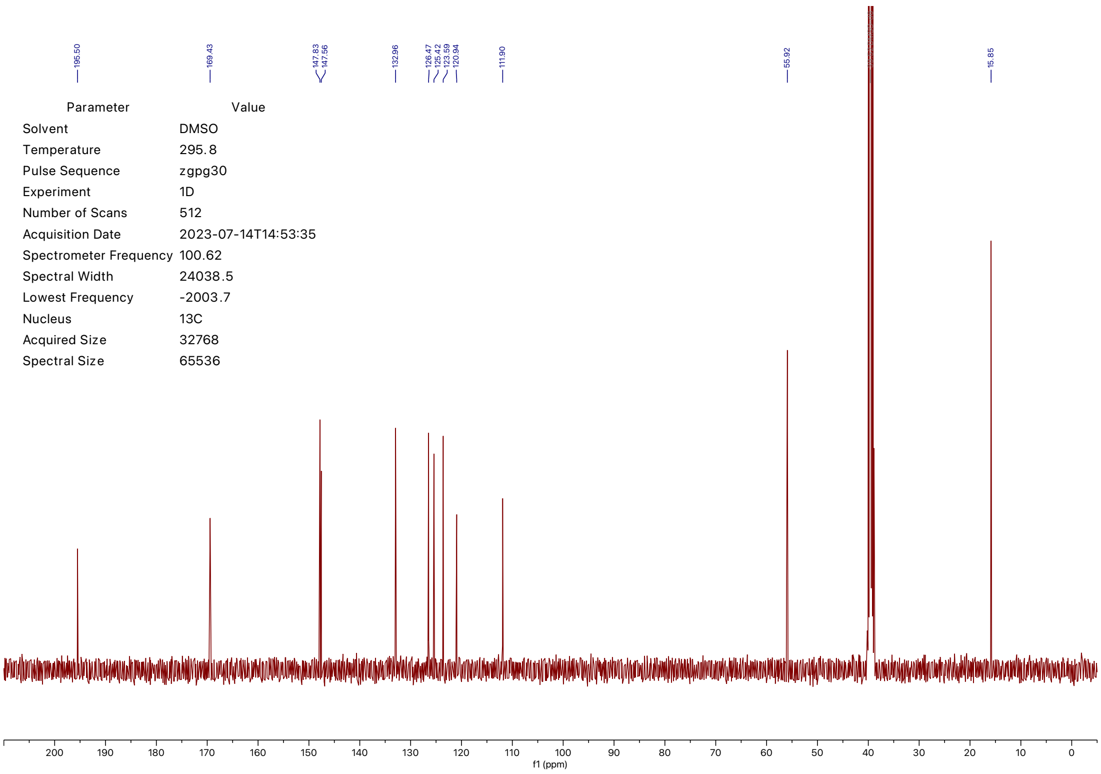


| Parameter | Value |
|---|---|
| Solvent | DMSO |
| Temperature | 295.8 |
| Pulse Sequence | zgpg30 |
| Experiment | 1D |
| Number of Scans | 512 |
| Acquisition Date | 2023-07-14T14:53:35 |
| Spectrometer Frequency | 100.62 |
| Spectral Width | 24038.5 |
| Lowest Frequency | -2003.7 |
| Nucleus | 13C |
| Acquired Size | 32768 |
| Spectral Size | 65536 |

| Parameter | Value |
| --- | --- |
| Solvent | DMSO |
| Temperature | 295.6 |
| Pulse Sequence | zg30 |
| Experiment | 1D |
| Number of Scans | 43 |
| Acquisition Date | 2021-04-10T18:34:00 |
| Spectrometer Frequency | 400.13 |
| Spectral Width | 8012.8 |
| Lowest Frequency | -1539.2 |
| Nucleus | 1H |
| Acquired Size | 32768 |
| Spectral Size | 65536 |

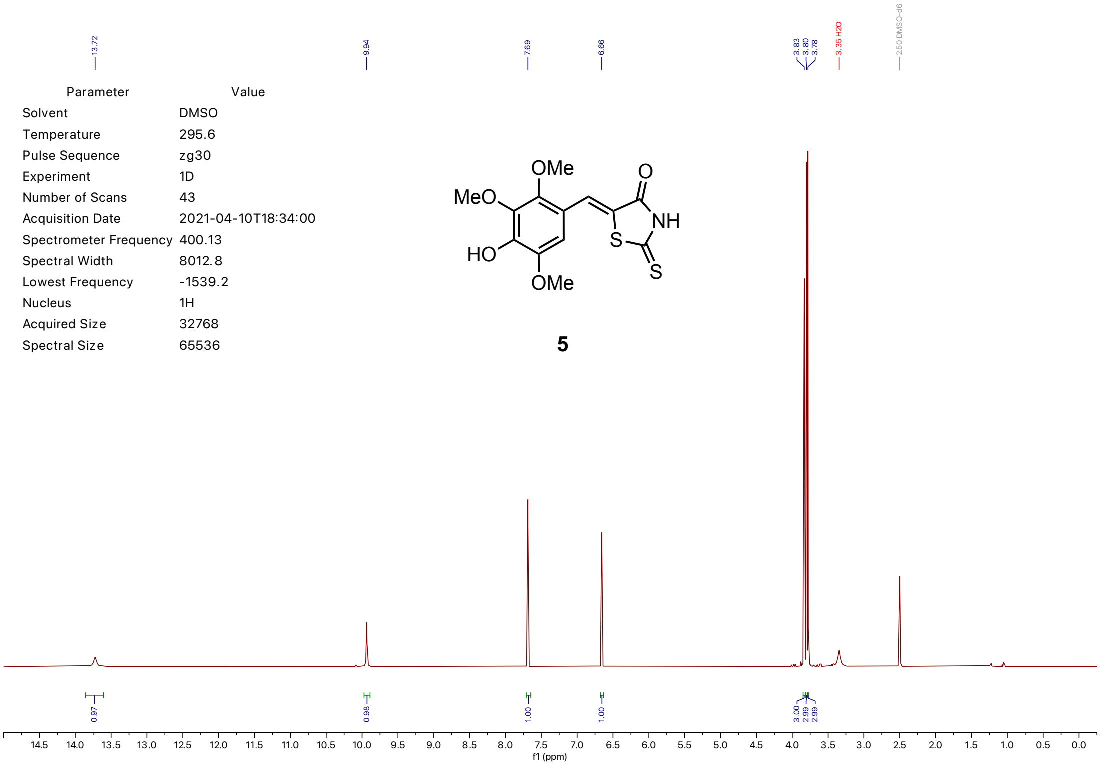


| Parameter | Value |
| --- | --- |
| Solvent | DMSO |
| Temperature | 296.0 |
| Pulse Sequence | zgpg30 |
| Experiment | 1D |
| Number of Scans | 20000 |
| Acquisition Date | 2021-04-10T18:41:00 |
| Spectrometer Frequency | 100.62 |
| Spectral Width | 24038.5 |
| Lowest Frequency | -2005.5 |
| Nucleus | 13C |
| Acquired Size | 32768 |
| Spectral Size | 65536 |

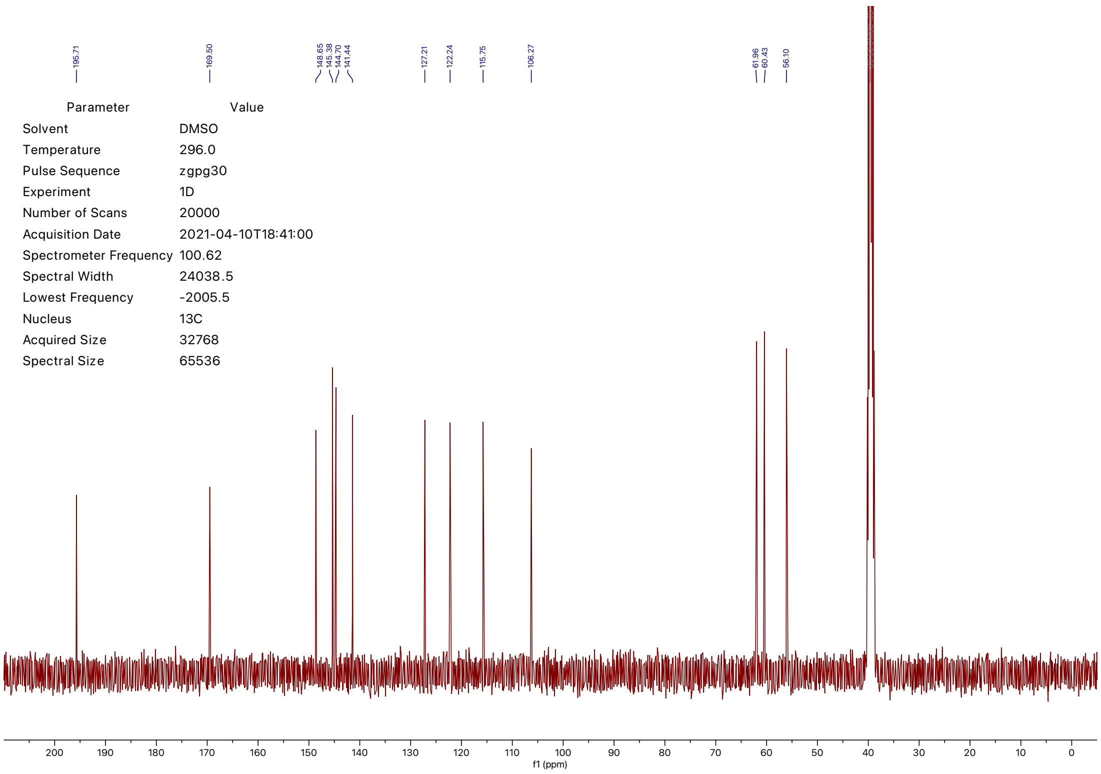

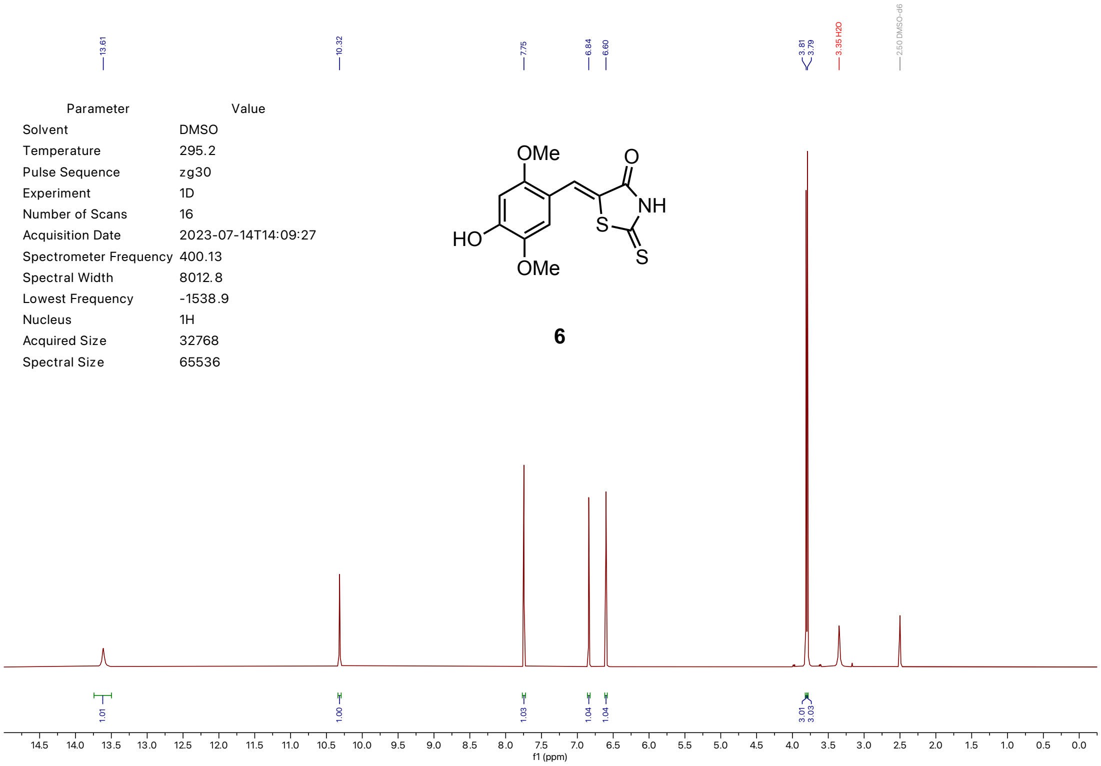


| Parameter | Value |
|---|---|
| Solvent | DMSO |
| Temperature | 295.2 |
| Pulse Sequence | zg30 |
| Experiment | 1D |
| Number of Scans | 16 |
| Acquisition Date | 2023-07-14T14:09:27 |
| Spectrometer Frequency | 400.13 |
| Spectral Width | 8012.8 |
| Lowest Frequency | -1538.9 |
| Nucleus | 1H |
| Acquired Size | 32768 |
| Spectral Size | 65536 |



**6**

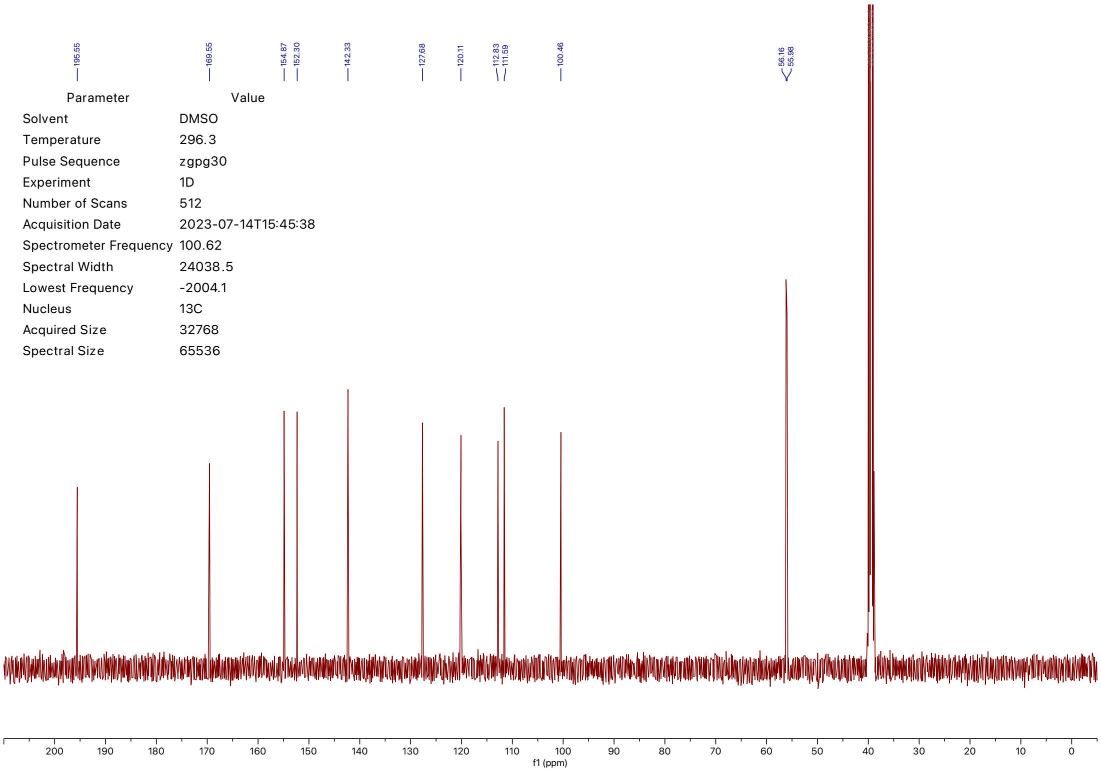


| Parameter | Value |
|---|---|
| Solvent | DMSO |
| Temperature | 296.3 |
| Pulse Sequence | zgpg30 |
| Experiment | 1D |
| Number of Scans | 512 |
| Acquisition Date | 2023-07-14T15:45:38 |
| Spectrometer Frequency | 100.62 |
| Spectral Width | 24038.5 |
| Lowest Frequency | -2004.1 |
| Nucleus | 13C |
| Acquired Size | 32768 |
| Spectral Size | 65536 |
